# Resolving the immune response across clonal cancer evolution in situ with Atera whole transcriptome profiling

**DOI:** 10.64898/2026.08.03.741871

**Authors:** Shaocheng Wu, Mohita Mahajan, Maarten van der Linde, ChunFang Zhu, Robert B West, David van IJzendoorn, Magdalena Matusiak

**Affiliations:** Department of Pathology, Stanford University, Stanford, USA; Department of Pathology, Leiden University Medical Center, Leiden, The Netherlands

**Author notes:** Corresponding authors: Correspondence to M.M.

## Abstract

Single-cell spatial transcriptomics is now central to studying tumors in their native tissue context. Here we present the first comprehensive, independent evaluation of Atera, a new spatial whole-transcriptome platform, compared with Xenium in human ductal carcinoma in situ (DCIS). We show that Atera enables granular cell-state annotation and resolves rare cell populations, experimentally validated by multiplex immunofluorescence (IF). We further show that its transcriptome-wide coverage enables inference of copy-number alterations at single-cell resolution, allowing us to reconstruct the clonal evolution of DCIS. We orthogonally confirm the inferred copy-number alterations by whole-genome sequencing of 16 microdissected tumor regions from consecutive tissue sections. Finally, by mapping the immune microenvironment onto this clonal architecture, we demonstrate the feasibility of tracking the changes in immune response along the clonal tumor evolution in situ. Together, our results establish Atera as a validated platform for tracking clonal evolution and immune adaptation in clinical samples.

## Main

Single-cell spatial transcriptome technologies can profile gene expression while preserving native tissue architecture. This allows cellular phenotype, cell state, and cellular organization to be defined in clinical patient samples^1,2^. Single-cell-resolution in situ platforms based on targeted gene panels (including MERFISH^3^, CosMx^2^, and Xenium^4^) have been adopted rapidly across the field. Yet the defining feature of these assays, a fixed panel of preselected genes, is also their central limitation: cell states not anticipated at panel design are difficult to resolve, rare populations defined by markers outside the panel may be missed, and genome-scale inferences such as somatic copy number variations (CNVs) remain limited. Although sequencing-based, spot-resolution spatial transcriptomics has been used to infer CNVs and tumor clonal evolution^5–12^, its multicellular resolution averages transcripts across neighboring cells and blurs clonal boundaries.

Whole-transcriptome in situ profiling has recently emerged to overcome the limitations of both existing approaches (the restricted gene coverage of targeted single-cell panels and the coarse, multicellular resolution of spot-based sequencing) by measuring the entire transcriptome at single-cell resolution in intact tissue. Early preprints applied a whole-transcriptome assay (WTA) on the CosMx platform (SMI®, Bruker) to reconstruct the normal-to-cancer transition in colon epithelium, ordering cells along a transcriptome-based trajectory^13,14^. These studies, however, inferred tumor evolution entirely from transcriptomic patterns, without validation against an independent, genomic ground truth.

Separately, several studies have benchmarked targeted imaging-based platforms against one another^4,15–19^, but these comparisons assessed agreement between platforms rather than validating their cell-type and cell-state calls against an independent measurement. Nor did they test how a platform’s sensitivity and panel size govern which rare cell populations and functional cell states can be recovered.

Two questions therefore remained unresolved: whether whole-transcriptome, single-cell profiling recovers rare cell states that targeted panels miss, and whether its genome-scale inferences hold up against direct DNA measurement. We address both with the first independent evaluation of Atera, a recently released spatial WTA from 10x Genomics, profiled alongside the Xenium v1 (280-probe) and Xenium Prime (5,000-probe) panels. Atera recovered functional cell states and rare cell populations beyond the reach of the targeted panels and enabled somatic copy-number inference at a granular resolution they cannot achieve. These capabilities extend in situ analysis from cell-state mapping to resolving tumor clonal architecture in tissue. To establish that this improved performance reflects biology rather than technical artifact, we employed two orthogonal experimental validations. First, we used multiplex immunofluorescence (IF) to confirm the presence and spatial distribution of rare cell populations. Second, we used whole-genome sequencing (WGS) of 16 different laser-capture-microdissected single-duct regions to validate the clonal tumor architecture inferred from the Atera WTA. To our knowledge, this is the first evaluation of copy-number calls from a single-cell spatial WTA against DNA sequencing. Applying this framework to DCIS, we mapped fine-grained immune annotations onto the tumor’s clonal architecture and traced how the immune response changes across clonal tumor evolution.

## Results

### Spatial mapping of DCIS and its tumor microenvironment

To compare Atera’s performance to targeted Xenium panels, we analyzed three publicly available 10x Genomics datasets of FFPE human breast tissue containing DCIS: the 280-gene Xenium v1 panel (Xenium 280), the 5,000-gene Xenium Prime panel with 100 custom genes (Xenium 5K), and a pre-production 18,000-gene Atera WTA. The Xenium 280 and Atera datasets were generated using sections from the same tissue block (∼100 µm apart), whereas the Xenium 5K dataset was derived from a different patient with a diagnosis of DCIS (Fig. 1a,b, first column). In addition to DCIS, the Xenium 5K section contains a region of invasive breast cancer (Supplementary Fig. 1a–c). Moreover, normal breast lobules and ducts were present in Atera and Xenium 280, but not in the Xenium 5K dataset.

**Fig. 1:**
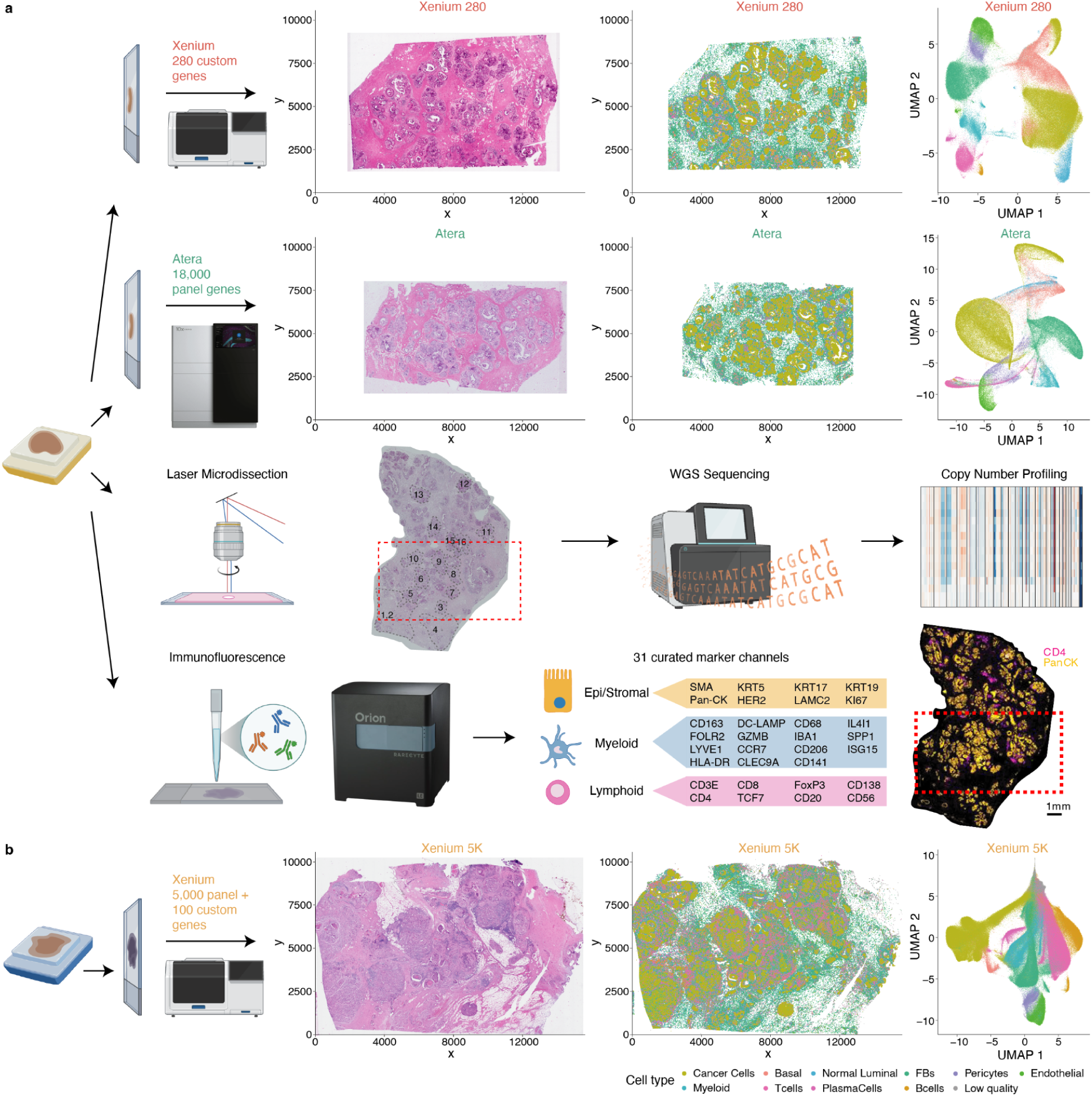
Study overview and cell-type annotation across the three spatial transcriptomics platforms. **(a)** Sections from the same FFPE human DCIS tissue block profiled with the 280-gene Xenium panel (Xenium 280), the 18,000-gene Atera WTA, laser-capture microdissection followed by whole-genome sequencing, and multiplex IF. The red box indicates a region of tissue that was profiled by Atera. **(b)** A second human DCIS tissue block was profiled with Xenium 5K (5,000-gene panel plus 100 custom genes). For each spatial transcriptomics platform, left to right: H&E image, spatial map, and UMAP colored by annotated cell types.

The diagnosis of DCIS was confirmed in the H&E staining of all datasets by an expert pathologist (Fig. 1a,b, second column). After quality filtering, library-size normalization and unsupervised clustering, we annotated the major cell types present in the tissue, including DCIS cancer cells (Cancer Cells), normal luminal cells (Normal Luminal), basal cells (Basal), fibroblasts (FBs), endothelial cells, pericytes, myeloid cells, T cells, B cells and plasma cells. Projecting these annotations back onto the tissue recovered the expected histological organization on every platform: cancer cells formed compact ductal structures embedded within a fibroblast-rich stromal and immune microenvironment (Fig. 1a,b, third column), and the corresponding UMAP embeddings resolved the same cell-type compartments in each dataset (Fig. 1a,b, fourth column). This shared cellular framework provided the common basis for the platform comparisons that follow.

### Comparative analysis of Xenium 280, Xenium 5K, and Atera

To compare the sensitivity of Xenium 280, Xenium 5K, and Atera, we analyzed the distribution of transcript counts and the number of genes detected per cell. Atera showed the highest transcript counts and gene detection per cell, followed by Xenium 280 and Xenium 5K (Fig. 2a,b). To further characterize the distribution of gene detection frequencies, we grouped genes by their average count per cell. We asked how many genes in the assay display only one or two counts per cell, and how this compares to genes with higher counts across the three platforms. Atera showed a higher number of genes across all transcript count bins compared with Xenium 280 and Xenium 5K (Fig. 2c). Next, we compared the spatial distribution of total transcript counts per cell (Fig. 2d), which showed the transcript density is highest in Atera, followed by Xenium 280 and Xenium 5K, consistent with our earlier comparisons (Fig. 2a).

**Fig. 2:**
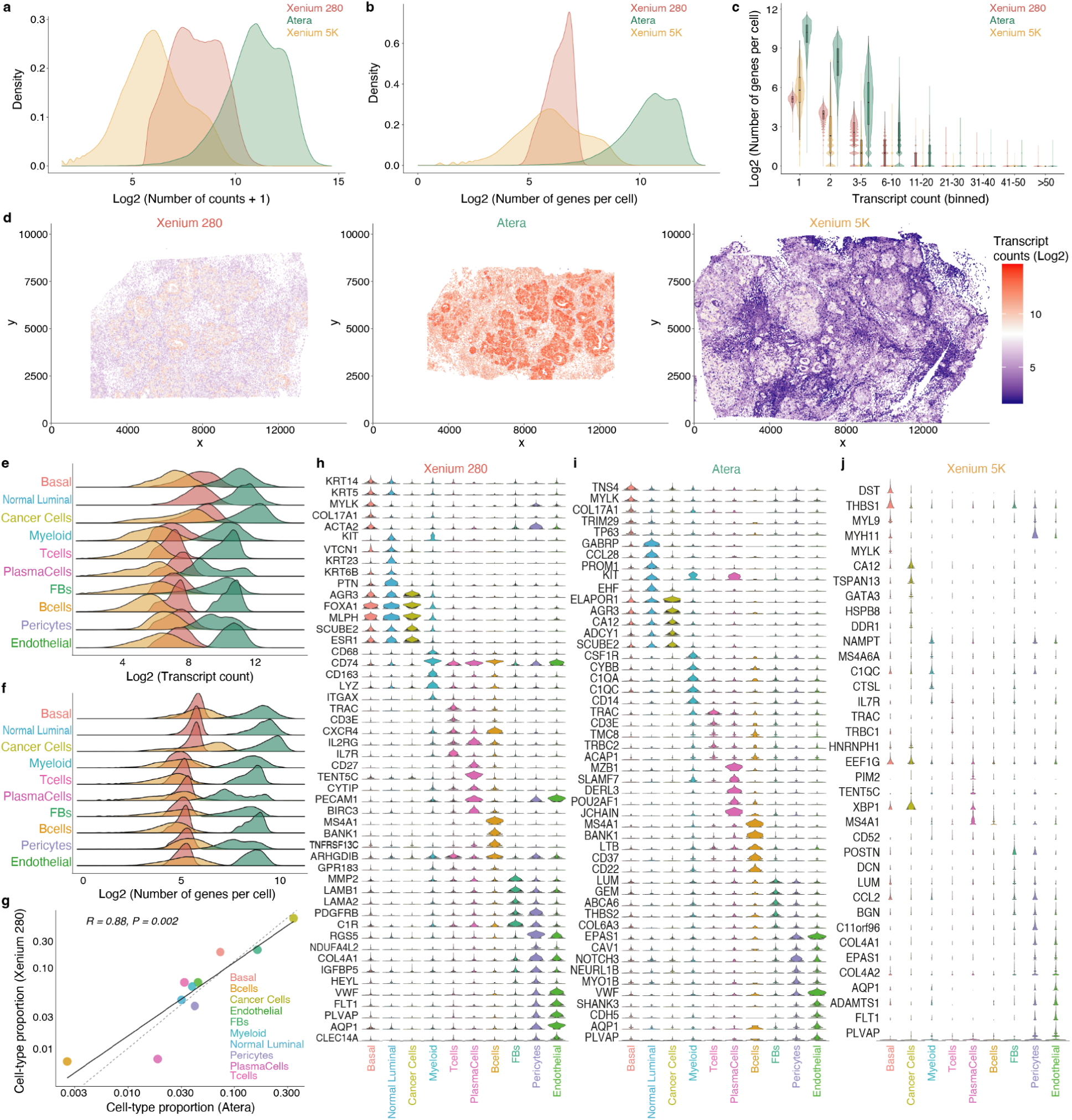
Performance comparison across Xenium 280, Xenium 5K, and Atera. **(a)** Density distribution of transcript counts per cell. **(b)** Number of genes detected per cell. **(c)** Violin plot showing the distribution of the number of genes detected per cell across different transcript count bins. Each bin represents a range of transcript counts detected within a cell. **(d)** Spatial distribution of the transcript count across the tissue. **(e)** Ridgeline plots showing the distribution of transcript counts per cell. **(f)** Number of genes detected per cell across the identified major cell types. **(g)** Scatter plot showing the correlation of cell type proportions between Xenium 280 and Atera. **(h–j)** Violin plots showing the distribution of transcript counts per cell for the top 5 differentially expressed marker genes of the corresponding cell types in Xenium 280, Atera, and Xenium 5K, respectively. The height of the violin plot shows the range of transcript counts for that gene across all the cells. Of note, the Xenium 5K sample did not contain normal breast ducts or lobules and thus, Normal Luminal cells are absent in the Xenium 5K dataset.

We next asked how these sensitivity differences affect the practical task of annotating cell types, which depends on detecting enough transcripts, genes and, most directly, cell-type marker genes in each cell. We therefore compared all three across the major cell types (Fig. 2e–j). Broken down by cell type, Atera detected higher transcript counts and more genes per cell than both Xenium panels across every major cell type (Fig. 2e,f); Xenium 280 detected more transcripts than Xenium 5K, although genes per cell varied by cell type.

Because Atera and Xenium 280 profiled consecutive sections of the same block, we compared the cell-type compositions they recovered; the proportions were strongly correlated (R = 0.88, P = 0.002; Fig. 2g), indicating that both platforms captured the same tissue composition despite their difference in sensitivity.

After that, we examined the signal most directly used for annotation: the expression of cell-type marker genes. For each platform, we identified the top five differentially expressed markers of each major cell type and compared their per-cell transcript counts across cell types (Fig. 2h–j). These cell type markers were most strongly and specifically expressed in Atera, intermediate in Xenium 280, and weakest in Xenium 5K. A marker set shared across all three platforms showed the same trend (Supplementary Fig. 2a).

Finally, to benchmark all three platforms against community quality standards, we evaluated the Xenium 280, Xenium 5K and Atera datasets with SpatialQM, a standardized quality-control framework for imaging-based spatial transcriptomics^20^. Sensitivity and coverage were highest for Atera: transcript density (transcripts per 100 µm² of cell area) was greatest for Atera and lowest for Xenium 5K (Supplementary Fig. 2b), and the per-gene count distribution, which reflects the dynamic range of quantification, was widest for Atera and narrowest for the 280-gene Xenium panel (Supplementary Fig. 2c). Atera also showed the most pronounced spatial structure, with the highest Moran’s I values across each platform’s top-50 expressed genes, followed by Xenium 280 and then Xenium 5K (Supplementary Fig. 2d).

Assignment efficiency and specificity were high on all three platforms. The fraction of transcripts assigned to cells (FTC), an index of segmentation quality, was high on all three (0.79, 0.89 and 0.80 for Xenium 280, Xenium 5K and Atera, respectively; Supplementary Fig. 2e). The signal-to-noise ratio (SNR), which indexes how far real signal rises above the negative-control background across the whole panel, was highest for Xenium 280 (2,154) and lower for Xenium 5K (688) and Atera (644). Because this metric averages signal over all panel genes, the broader 5,000-gene and whole-transcriptome (∼18,000-gene) panels, which include many lowly expressed genes, are diluted relative to the focused 280-gene panel, so their lower values reflect panel breadth rather than greater noise (Supplementary Fig. 2f). The corresponding negative-control-derived false-discovery rate (FDR = 1/SNR) followed the same pattern, lowest for Xenium 280 and highest for Atera (0.046%, 0.145% and 0.155% for Xenium 280, Xenium 5K and Atera) (Supplementary Fig. 2g). Finally, the mutually-exclusive correlation (MECR, the per-cell correlation between markers of different lineages) was negative for spatially distinct (primary) lineage pairs on every platform (−0.13, −0.01 and −0.08), confirming clean segregation of cell types that occupy separate locations, while co-localizing (secondary) pairs were positive, as expected for cell types that are spatially co-enriched (Supplementary Fig. 2h).

Like SNR, sparsity and entropy were shaped by panel composition, not data quality. Sparsity (the fraction of panel genes with zero counts) was highest for the shallowly sampled Xenium 5K panel (0.98), intermediate for Atera (0.90), and lowest for Xenium 280 (0.79) (Supplementary Fig. 2i), while entropy (the evenness of expression across genes) rose with panel breadth, reaching a maximum of 12.96 bits for Atera (Supplementary Fig. 2j).

### Luminal DCIS clones are segregated in space

Clustering the Atera luminal epithelium resolved four transcriptional DCIS states (Luminal DCIS1 - 4) alongside the normal luminal state (Fig. 3a). The four epithelial states were enriched across spatially segregated territories in the tissue (Fig. 3b). We wondered whether these distinct regions could correspond to different tumor clones. To evaluate the feasibility of using Atera spatial transcriptomic data to infer genomic variation in the tumor compartment, we first asked whether the gene content of the spatial assay was sufficient to infer CNVs. The 280-and 5,000-gene Xenium panels cover only ∼1% and ∼25% of the human protein-coding genes, respectively, whereas the Atera WTA captures around 90%. Consequently, most chromosome arms carried only from 10 to 130 genes on the Xenium panels, versus around 440 genes per arm for Atera (Supplementary Fig. 3a).

**Fig. 3:**
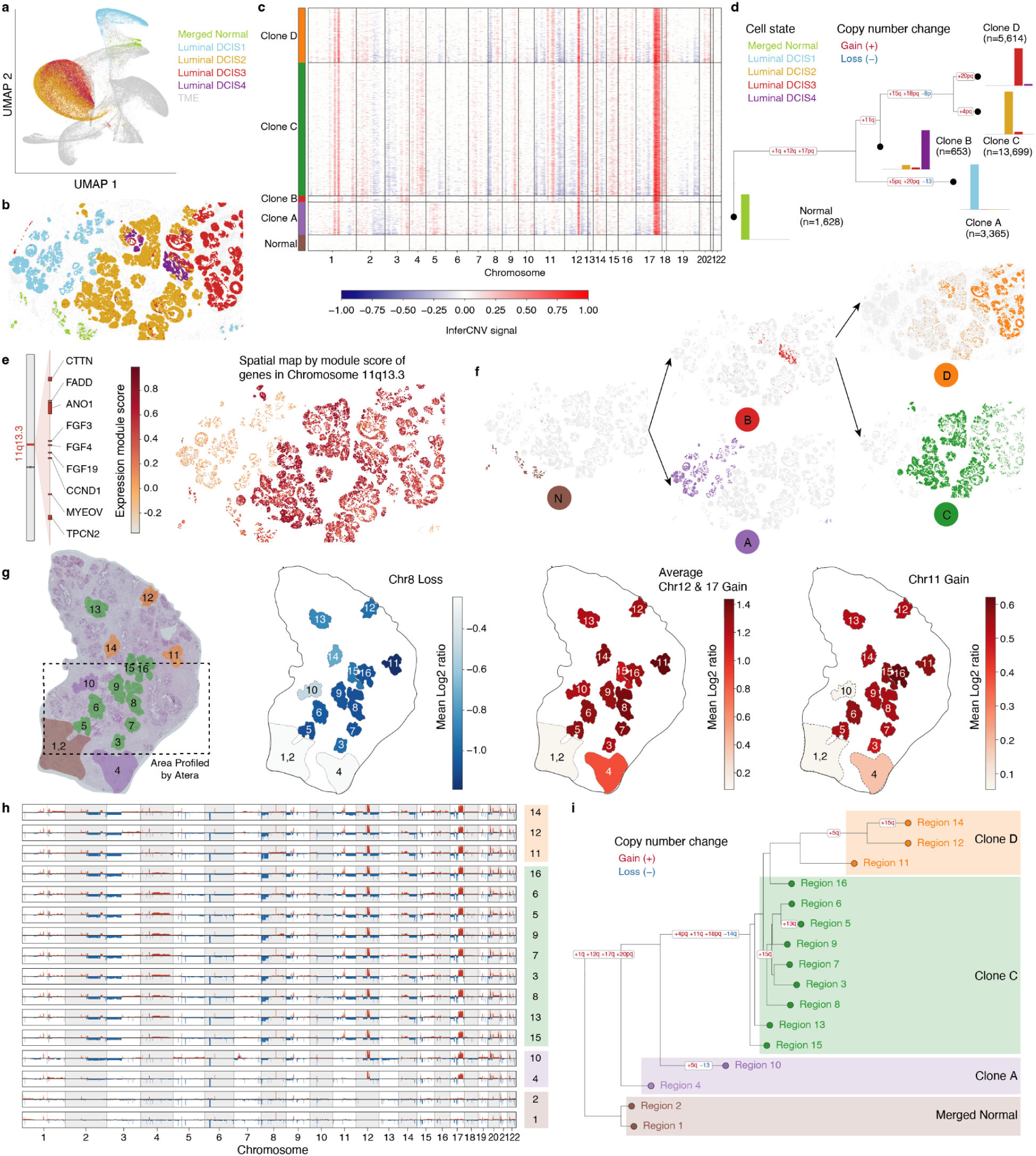
Resolving clonal evolution of DCIS by copy number inference using Atera. **(a)** UMAP colored by luminal DCIS states. **(b)** Spatial map of the luminal DCIS states. **(c)** Single-cell inferCNV heatmap of the Atera DCIS epithelium (cells grouped by inferCNV clone). **(d)** Phylogenetic tree of the clone-level CNV profiles, rooted on Merged Normal. **(e)** Spatial map of the 11q13.3 amplicon module score (CCND1, FGF3/4/19, ANO1, CTTN, FADD, MYEOV, TPCN2). **(f)** Spatial clonal evolution of the luminal DCIS clones. **(g)** H&E with the 16 laser-microdissected regions colored by CNV clones, and spatial maps of mean log2 ratio for chr8 loss, average chr12/17 gain, and chr11 gain. **(h)** WGS copy-number profiles for the 16 laser-microdissected regions. **(i)** Phylogenetic tree of the region-level WGS profiles, rooted on the near-diploid regions.

We first ran inferCNV on the targeted Xenium 5K and 280 datasets, using normal luminal or myoepithelial cells as the reference. Although both panels detected chromosome-scale gains and losses that separated the DCIS epithelium from the reference (Supplementary Fig. 3b and 3c), the 280-gene panel carried insufficient genomic coverage for reliable copy-number inference. The inferred resolution scaled directly with panel breadth: the 280-gene panel resolved only ∼22 genomic bins (approximately one per chromosome), restricting it to whole-chromosome events, whereas the 5,000-gene panel resolved approximately 280 bins and recovered arm-level alterations (gains on chr1, chr8, chr11 and chr17). With the broadest coverage, the Atera WTA generated ∼1,500 bins. Correlation against the matched WGS ground truth confirmed this limitation: the Atera copy-number profiles were highly concordant with the DNA-based calls, whereas the Xenium 280 profiles were not (Supplementary Fig. 3d).

In an independent breast tumor profiled with the Xenium 5K panel, the arm-level calls recovered known recurrent breast cancer CNVs^21–25^: 1q gain with 16q loss, 8q (MYC) gain with 8p loss, 17q (ERBB2) gain with 17p (TP53) loss, and gains at 11q13 (CCND1), suggesting that a sufficiently broad targeted panel could capture genuine copy-number biology.

Applying inferCNV to the Atera epithelial compartment with normal and ductal luminal cells as reference, we recovered a clear CNV landscape that was absent from the normal reference cells (Fig. 3c). Unsupervised clustering of the per-cell CNV profiles partitioned the luminal DCIS epithelium into four discrete CNV clones spanning from a dominant clone (Clone C, n = 13,699 cells) through intermediate clones (Clone D, n = 5,614; Clone A, n = 3,365) to a minor subclone (Clone B, n = 653), against a Normal reference (n = 1,628) (Fig. 3c)

Next, we sought to reconstruct a phylogenetic tree using Atera’s clonal CNV profiles to order the DCIS tumor clones into a branching evolutionary hierarchy (Fig. 3d). A set of truncal gains (+1q, +12q, +17pq) was shared by all tumor clones, defining the common malignant founder. After these common events, the clones diverged through clone-specific events: Clone A branched early via a distinct trajectory (+5pq, +20pq, −13), whereas the remaining clones shared a subsequent gain of chromosome 11 (+11q: 12.1-13.1 and 13.3-14.1) starting with Clone B and then further diversified along an internal branch (+15q, +18pq, −8p) before splitting into Clone C (+4pq) and Clone D (+20pq). This establishes a normal-to-tumor progression captured entirely within a single tissue section (Fig. 3d).

A central observation from this hierarchy is that the chromosome 11q gain was not a truncal event but was acquired on an internal branch. This marks the lineage leading to Clones B, C, and D (Fig. 3d). This 11q gain was absent from the early-diverging Clone A. Because Atera retains the spatial coordinates of every cell, this clonal distinction is translated into a spatial one. Mapping an 11q13.3 amplicon-gene expression module (CCND1, FGF3/4/19, ANO1, CTTN, FADD, MYEOV, TPCN2) onto the spatial tissue architecture revealed pronounced spatial heterogeneity, with amplicon-high and amplicon-low territories occupying distinct spatial regions rather than being uniformly distributed (Fig. 3e). This expression pattern was concordant with the underlying inferred 11q copy-number signal. Consistent with this clonal architecture, the four Luminal DCIS clones and the Normal reference occupied distinct, spatially segregated territories across the tissue (Fig. 3f).

To validate these transcriptome-based clonal CNV calls with an orthogonal DNA-based measurement, we laser-microdissected 16 spatially distinct DCIS tumor ducts across two sections from the same tissue block as profiled by Atera and Xenium 280, and profiled them by WGS (Fig. 1a). Mapping the WGS-derived copy-number ratios back onto the physical location of each dissected region reproduced the spatial organization seen in the transcriptome-based clones: the chromosome 8 loss and the chromosome 12 and 17 gains were similarly regionally confined and regions carrying the chromosome 11 gain were spatially segregated from the one lacking it (region 10) (Fig. 3g). The region-level WGS copy number profiles recovered the same landmark alterations inferred from the Atera transcriptome (including the chromosome 1, 11, 12, and 17 gains and the chromosome 8 loss) across the microdissected regions (Fig. 3h and Supplementary Fig. 3e). Phylogenetic tree reconstruction of these region-level profiles grouped the dissected regions into clones corresponding to the clones defined by the Atera transcriptome (Clones A, C and D), rooted on the normal reference regions (Fig. 3i). This orthogonal, DNA-level confirmation validates the spatially segregated clonal architecture inferred from the Atera WTA assay.

### Resolving myeloid cell types and states

To test the ability of the three platforms to resolve rare cell types and granular cell states we subseted and independently reclustered the myeloid compartment from tumors profiled with Xenium 280, Xenium 5K and Atera. Across all three cluster-quality metrics (silhouette, cLISI and 1 − kBET acceptance), myeloid populations were best resolved in Atera, followed by Xenium 5K and then Xenium 280 (Supplementary Fig. 4a).

Consistent with the cluster-quality metrics, the platforms differed in the granularity of the myeloid cell states they resolved. Atera resolved the most states, with well-separated clusters of classical and non-classical monocytes, FOLR2 tissue-resident macrophages (FOLR2 TRMs), SPP1 and CXCL9 tumor-associated macrophages (SPP1 TAMs and CXCL9 TAMs), and type 1 and type 2 conventional dendritic cells (cDC1s and cDC2s) (Fig. 4a). Atera alone also resolved the rare mature regulatory DCs (mregDCs) and plasmacytoid DCs (pDCs), which formed small, well-separated satellite clusters (Fig. 4a). We validated the presence of pDCs by Orion multiplex immunofluorescence on a tissue section adjacent to the one profiled by Atera (Supplementary Fig. 4b). By contrast, Xenium 5K recovered most of the common populations but did not resolve non-classical monocytes, mregDCs or pDCs, and it yielded a large C1QC-expressing cluster of likely macrophage identity that could not be assigned to a defined state, consistent with the higher marker-gene dropout on this platform relative to Xenium 280 and Atera (Fig. 4b). Xenium 280 resolved the fewest, just three states (classical monocytes, FOLR2 TRMs, and SPP1 TAMs), alongside monocyte and DC clusters that could not be confidently assigned (Fig. 4c,d).

**Fig. 4:**
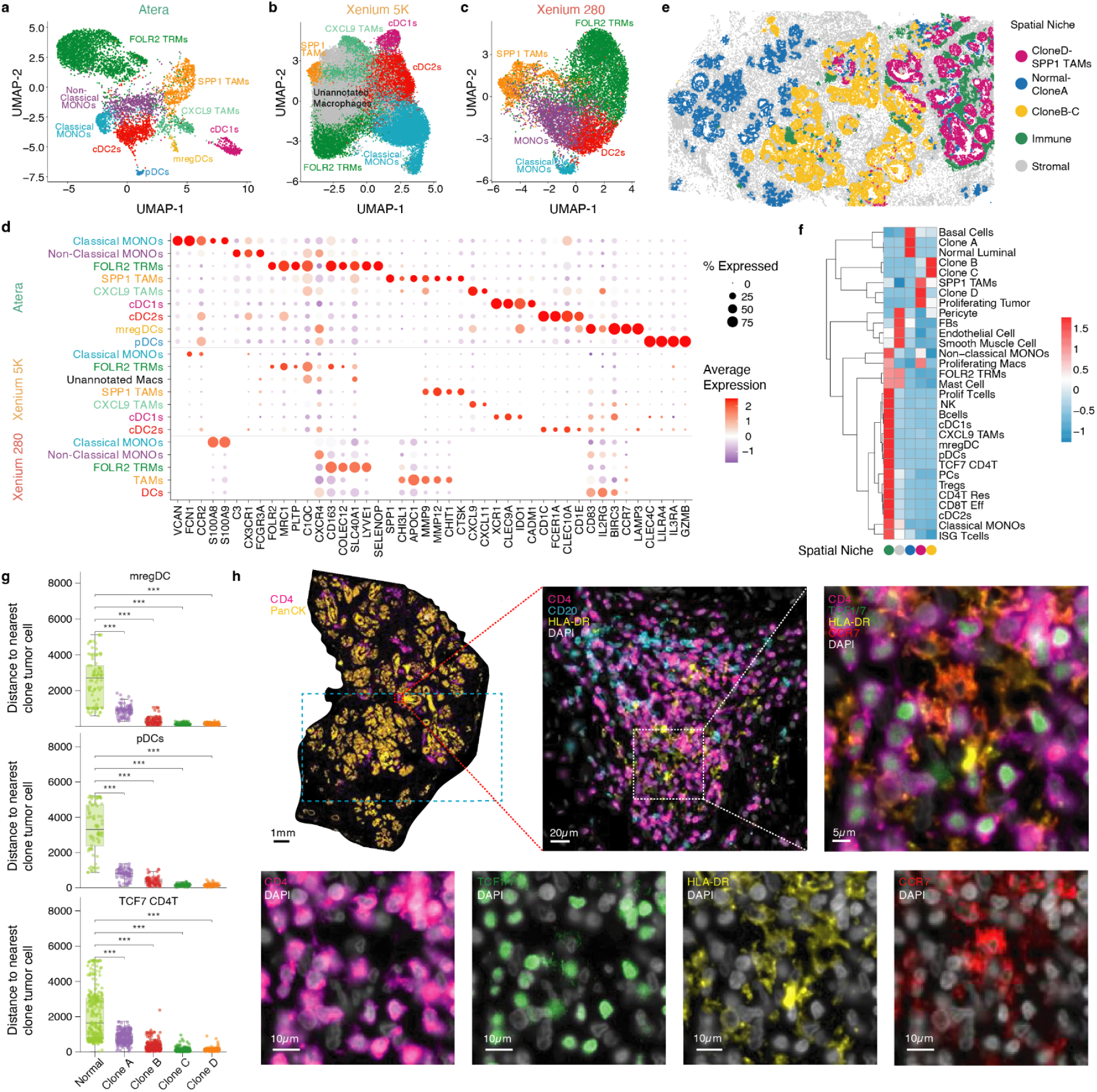
Resolving rare myeloid and immune cell states and their spatial organization around DCIS clones. **(a–c)** UMAP projections of reclustered myeloid cells from Atera **(a)**, Xenium 5K **(b),** and Xenium 280 **(c)**. **(d)** Dotplot of average gene expression per myeloid cluster across Atera, Xenium 5K and Xenium 280. **(e)** Spatial map of the five cellular niches across the Atera DCIS section. **(f)** Heatmap of the cell-type composition of each spatial niche. **(g)** Distance from mregDCs (top), pDCs (middle) and TCF7^+^ CD4 T cells (bottom) to the nearest clonal tumor cell and normal cell. **(h)** Orion multiplex IF on a tissue section adjacent to the one profiled by Atera, shown as a whole-section overview (CD4, Pan-CK), a magnified view showing co-localization of dendritic cells (HLA-DR), CD4+ T-cells (CD4) and B-cells (CD20). A further inset shows TCF1/7 positive CD4+ T-cells (TCF1/7 and CD4) and mregDCs (HLA-DR and CCR7), and single-marker panels for CD4, TCF1/7, HLA-DR and CCR7 (with DAPI).

To identify the technical basis for these differences in myeloid-state resolution, we subsetted the raw Atera counts by reducing either the number of genes or transcript detection depth (Supplementary Fig. 4c). Cluster separability (cLISI and 1 − kBET acceptance) was largely preserved at the 5,000-gene set, declined modestly at the 280-gene set, and fell most sharply when detection depth was reduced. Both the number of genes in the assay and the number of detected transcripts per cell, therefore, limit myeloid cell state recovery, with detection depth as the dominant factor.

### Tracking changing immune response along clonal DCIS evolution

We next examined how all tissue cells were spatially organized relative to the DCIS clones. Partitioning the Atera DCIS section into five recurrent spatial niches, defined by local cell-type composition, revealed an immune-enriched niche, a stromal niche, and three epithelial niches, each dominated by different DCIS clones (Fig. 4e,f). Normal luminal and Clone A cells resided almost exclusively in one niche (100% and 97.9%), Clones B and C were co-enriched in a second (84.4% and 88.4%), and Clone D occupied a third (81.7%). The rare mregDC, pDC and TCF7^+^ CD4 T cell populations mapped to the immune niche (Fig. 4f) and were positioned significantly closer to the DCIS tumor clones than to normal luminal epithelium (Fig. 4g). We confirmed this arrangement at the protein level by Orion multiplex immunofluorescence on a tissue section adjacent to the one profiled by Atera where TCF7^+^ CD4+ T cells and HLA-DR^+^ CCR7^+^ mature dendritic cells co-localized within tumor-adjacent immune aggregates (Fig. 4h).

Next, we sought to quantify the spatial association of the immune cells with DCIS tumor clones. To do that, we grouped DCIS tumor cells into ducts (Fig. 5a,b) and quantified the proportion of the immune niche within the 200 µm radius surrounding each DCIS duct (Fig. 5c). This revealed that the fraction of immune niche surrounding DCIS ducts increased between DCIS clones A, C, and D, and was consistently high across Clone D ducts. Because clonal stage is an ordered variable, we tested this trend with the Jonckheere–Terpstra ordered-trend statistic against a torus-shift permutation null, which preserves the spatial structure of the immune-niche field, and found it significant (JT z = 7.43, one-sided P = 0.004, 999 permutations; Fig. 5d). In addition, this analysis showed that the fraction of the immune niche increases along the x axis of the tissue from left to right (Fig. 5e). The smallest of all clones, Clone B, whose ducts are located between the spatial territories occupied by Clones C and D, was an exception to this trend.

**Fig. 5:**
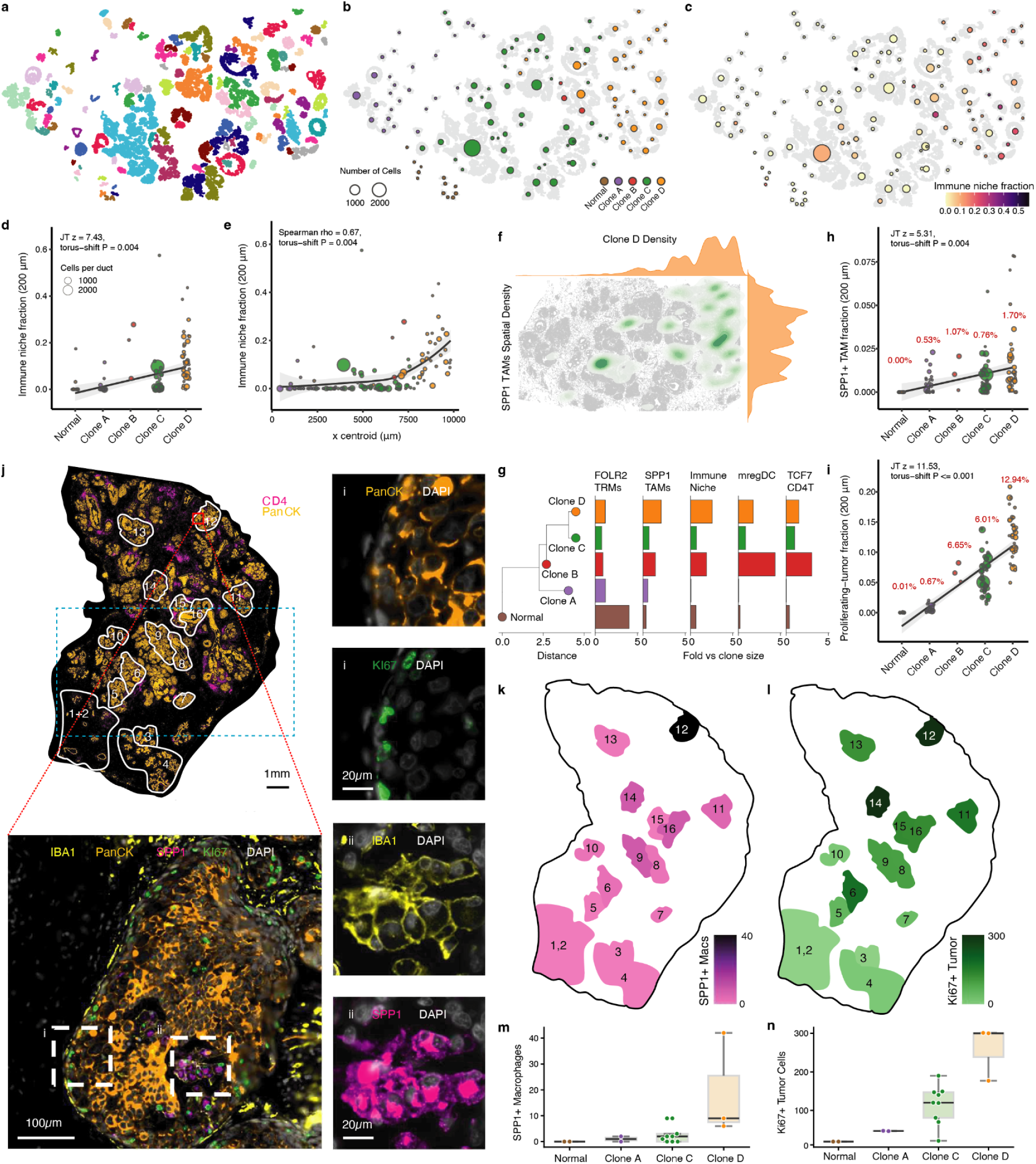
The immune response and luminal DCIS clones. **(a)** Duct segmentation of the tumor compartment by single-linkage clustering at 40 µm; colors denote duct identity (n = 128 ducts, 21,587 cells). **(b)** Duct centroids colored by their majority clone (Normal and Clones A–D; point size denotes the number of cells per duct). **(c)** Duct centroids colored by the immune-niche fraction within 200 µm of the duct. **(d)** Immune-niche fraction within 200 µm for each group (point size denotes cells per duct; JT z = 7.43, torus-shift P = 0.004). **(e)** Immune-niche fraction within 200 µm versus duct position along the spatial x axis (Spearman ρ = 0.67; torus-shift P = 0.004). **(f)** Joint spatial density of SPP1 TAMs (green) and Clone D tumor cells (orange) across the Atera section, with marginal distributions. **(g)** Fold enrichment of the immune niche and of FOLR2 TRMs, SPP1 TAMs, mregDCs and TCF7+ CD4 T cells near each clone, ordered along the clonal phylogeny (calculation included the proliferating tumor cells near each clone). **(h)** SPP1+ TAM fraction within 200 µm (per-clone mean fraction values are shown in red). **(i)** Proliferating-tumor fraction within 200 µm (per-clone mean fraction values are shown in red). **(j)** Orion multiplex IF (IBA1, PanCK, SPP1, Ki67, DAPI selected) on the adjacent tissue section (Top left: whole-section overview with the 16 laser-microdissected WGS regions; Bottom left: a highlighted region with single marker insets (i, ii) on the right showing proliferating tumor cells (PanCK+/Ki67+) and SPP1 TAMs (IBA1+/SPP1+) co-localized in the Clone D epithelium). **(k)** Spatial map of SPP1+ macrophage abundance across the 16 microdissected regions. **(l)** Spatial map of Ki67+ tumor-cell abundance across the same regions. **(m)** SPP1+ macrophages per region grouped by Normal and clones. **(n)** Ki67+ tumor cells per region grouped by Normal and clones.

Beyond the immune niche present in the stroma surrounding the DCIS ducts, SPP1 TAMs and proliferating tumor cells were spatially co-enriched within the epithelial compartment of Clone D (Fig. 4f and 5f,g). This finding is consistent with our previous report that SPP1 TAMs surround and infiltrate tumor islands while being absent from normal adjacent stroma^26^. We found that, SPP1^+^ TAM density rose from 0.53% within Clone A ducts to 1.70% within Clone D (P = 0.004; Fig. 5h), and the proliferating tumor fraction rose from 0.67% to 12.94% over the same span (P ≤ 0.001; Fig. 5i). We confirmed this co-localization at the protein level by Orion multiplex IF on a section adjacent to the one profiled by Atera and WGS (Fig. 5j). Across ducts corresponding to the 16 WGS-profiled regions, both SPP1^+^ macrophages and proliferating tumor cells increased with clonal advancement and were most abundant around Clone D (Fig. 5k–n).

## Discussion

In this study, we present an integrated framework for resolving the immune response across clonal cancer evolution within intact tissue. We provide the first evaluation of the Atera WTA and compare it to the targeted Xenium v1 and Xenium Prime panels. Within a single tissue section, the framework unifies single-cell copy-number inference, fine-grained annotation of immune cell states, and the spatial co-registration of both, enabling us to partition the DCIS epithelium into spatially resolved clones, reconstruct their phylogenetic relationships, and position the immune response along the resulting evolutionary hierarchy.

A defining feature of this work is that it uses adjacent sections from a single tissue block to experimentally validate findings derived from Atera WTA profiling. Prior comparisons of spatial platforms have benchmarked assays against one another to establish internal consistency. We extend this approach by assessing how faithfully the measurements reflect the underlying biology, using two orthogonal sources of ground truth. First, we used multiplex IF to confirm, at the protein level, the presence of the rare immune populations and their spatial organization around the tumor. Second, we used WGS to validate the inferred clonal architecture. To our knowledge, copy-number calls derived from a single-cell spatial WTA have not previously been evaluated against DNA sequencing. Using this approach, we show that CNVs as small as 6.7 megabases long at 11q12–q13.1 can be reliably detected. Together, these independent measurements substantiate the biological accuracy of the results inferred based on the Atera WTA.

In addition, previous comparisons of spatial platforms have focused largely on sensitivity, specificity, and signal contamination, but have not asked how panel breadth and assay depth govern the ability to resolve fine functional cell states and rare populations. Our findings indicate that both panel breadth and sensitivity influence the assay’s ability to recover rare cell types and resolve functional cell states. Supporting the importance of the number of profiled genes, we found that the 280-gene Xenium v1 panel resolved only three confidently assignable myeloid cell states, whereas the Atera WTA, which profiled a nearby tissue section from the same block, recovered a substantially broader repertoire of myeloid cell types and states. This includes rare cell populations (mregDCs and pDCs) and granular functional cell state annotation (recovery of CXCL9 TAMs and Non-Classical Monocytes). Conversely, the importance of the number of detected genes per cell was supported by a large cluster of myeloid cells that could not be confidently annotated due to the absence of cell state-defining markers in the 5,000-gene Xenium Prime panel.

Jointly resolving the genomic and immune axes within the same tissue exposes organization that is not apparent from either axis in isolation. In DCIS, whether a lesion remains indolent or progresses to invasive disease is thought to be shaped by the reciprocal interactions between genomically distinct subclones and the surrounding immune response^27,28^, an interplay that has been difficult to examine because it requires both the genome and the microenvironment to be read in the same section. More broadly, the ability to track the immune response to evolving tumor clones could help clarify how immune pressure shapes, and is shaped by, clonal selection across all tumor types.

Several limitations should be noted. Our clonal and immune analyses with Atera are derived from a single DCIS specimen, and the specific evolutionary trajectory and immune associations reported here will require validation in larger cohorts. Therefore, this study is best regarded as an illustration of the biology that this approach can access rather than a population-level generalization. We benchmarked Atera against Xenium panels because (1) Xenium is one of the most widely adopted single-cell in situ platforms¹⁷,¹⁸, (2) it is produced by the same vendor, and (3) a publicly available Xenium v1 dataset had been generated from the same tissue block. Extending the comparison to spatial WTA platforms from other vendors, such as CosMx (Bruker), will be an important next step. Notwithstanding these caveats, our study presents an orthogonally validated framework to link clonal cancer evolution and the immune response within intact tissue using a single-cell spatial WTA.

## Author Contributions

S. Wu and M. Matusiak conceived this study; M. van der Linde and C. Zhu performed experiments; S. Wu, M. Mahajan, M. van der Linde and D. van IJzendoorn analyzed the data; R.B. West provided histology expertise; R.B. West, D. van IJzendoorn and M. Matusiak provided resources; S. Wu, M. Mahajan, M. van der Linde and M. Matusiak wrote the original draft; all authors contributed to manuscript review and editing; R.B. West and M. Matusiak acquired funding for this study.

## Acknowledgments

This work was supported by grants from the National Cancer Institute R01CA229529 to R.B. West and M. Matusiak, and R01CA290021 to R.B. West. We did not receive financial or editorial support from 10x Genomics. We thank Syrus Mohabbat (10x Genomics) for providing the tissue sections adjacent to the sections used for Xenium v1 and Atera profiling and Amanda Janesick (10x Genomics) for orienting the Xenium v1 and Atera sections relative to those shared with our lab. During the preparation of this work, the authors used Claude to assist with rewriting the manuscript text and R code for data analysis and visualization. The authors reviewed, edited, tested, and validated all Claude outputs.

## Methods

### Data sources

All spatial single-cell datasets used in this study are available from the 10x Genomics website. Specifically, Xenium 280 data was obtained from the S1-Bottom section in https://www.10xgenomics.com/datasets/xenium-ffpe-human-breast-biomarkers. Pre-production Atera Whole Transcriptome Assay data was obtained from https://www.10xgenomics.com/datasets/atera-wta-ffpe-human-breast-cancer. Xenium 5K data was obtained from https://www.10xgenomics.com/datasets/xenium-prime-ffpe-human-breast-cancer.

### Data preprocessing and cell type annotation

For all 3 datasets, the cell-by-gene raw count matrix, per-cell metadata, and run metadata were imported into R and assembled into a Seurat object (Seurat v4; R v4.2). The raw count matrix was filtered such that each cell had at least 50 transcript counts, and then they were library-size normalized and log-transformed. Highly variable features were identified, and the expression matrix was scaled before principal-component analysis (PCA). A nearest-neighbor graph was built on the top 30 principal components, and a UMAP embedding was computed for visualization. Cells were then partitioned with the Leiden algorithm. For each cluster, the top 30 differentially expressed marker genes were identified and retained for annotation. Manual broad cell type annotation was performed based on the top 30 differentially expressed marker genes and cell types’ canonical marker genes.

### Sensitivity comparison among Xenium 280, Xenium 5K, and Atera

Total transcript counts and the number of genes detected per cell were compared across Xenium 280, Xenium 5K, and Atera. Low-quality cells were removed, and per-cell transcript counts and gene counts were log2-transformed and displayed as density curves, both overall and within each major cell type. Spatial distribution was visualized by coloring each cell at its tissue coordinates by log2-transformed transcript count, using a common color scale across platforms with limits set to the 5th and 95th percentiles. For Fig. 2c, Genes were grouped per cell into transcript-count bins (1, 2, 3–5, 6–10, 11–20, 21–30, 31–40, 41–50 and >50 transcripts), and the number of genes per bin was counted, log2-transformed, and shown as violin plots with boxplots. For each major cell type, the per-cell transcript counts of its top 5 differentially expressed marker genes were log2-transformed and displayed as violin plots. For gene-level sensitivity, overlapping genes were identified for each platform pair, the mean transcript count per cell was computed for each gene, and the two platforms were compared on log10-scaled scatter plots. For Xenium 280 and Atera, which were profiled from nearby sections of the same tissue block, major cell-type proportions were computed per platform and compared by Spearman correlation.

### SpatialQM metrics calculation

SpatialQM metrics were computed for each dataset. Transcript density was the median of 100 times (per-cell gene counts / cell area in µm²). Dynamic range was the base-10 log ratio of the highest- to lowest-expressed gene (total counts per gene across cells). Spatial autocorrelation was the median per-gene Moran’s I of the top50 most-expressed genes. The fraction of transcripts in cells (FTC) was the sum of in-cell gene counts divided by total high-quality decoded transcripts. Signal-to-noise ratio (SNR) was the mean counts per gene divided by the mean counts per Negative Control Probe. SpecificityFDR was calculated as the reciprocal of SNR. The mutually-exclusive correlation (MECR) was quantified as the mean across-cell Pearson correlation of the canonical marker pairs of mutually exclusive lineages, reported separately for spatially distinct (“primary”) pairs (EPCAM–CD3E, EPCAM–MS4A1, EPCAM–CD68, EPCAM–PECAM1, EPCAM–PDGFRB, CDH3–CD3E, and CDH3–PDGFRB) and co-localizing (“secondary”) pairs (CD3E–MS4A1, CD3E–CD68, CD68–MS4A1, and PECAM1–PDGFRB). Per-dataset sparsity was the fraction of zero entries in the cell-by-gene matrix. Entropy was calculated as the Shannon entropy of the aggregate gene-expression distribution (H = −Σ p_g log₂ p_g).

### Atera data copy number inference

Single-cell copy-number profiles were inferred with inferCNVpy (v0.6.1) across the complete Atera dataset, using Normal Luminal and Ductal Luminal cells as the reference and all remaining tumor cells as observations. Genes were assigned GRCh38 chromosomal coordinates by symbol, ordered by genomic position and restricted to autosomes (chr1–chr22), yielding 17,237 ordered genes, and per-cell genome-wide profiles were estimated over a sliding 100-gene window, centered and de-noised relative to the reference mean. To define copy-number clones on high-quality cells while preserving the diploid baseline, the per-cell CNV profiles were then retained unchanged (no re-inference) and a gene-detection filter was applied to the observation cells only: observation cells with fewer than 3,000 detected genes were removed, whereas all reference cells were retained regardless of gene count. On this filtered set, principal component analysis, a nearest neighbor graph and Leiden clustering were recomputed to obtain fresh cluster assignments, and clusters of fewer than 260 cells were merged into a single catch-all group. The resulting clusters were used as the CNV clones for all downstream analysis.

### Laser capture microdissection

Unstained sequential sections from the DCIS FFPE block profiled with Atera were provided by 10x Genomics on Epredia^TM^ Superfrost^TM^ Plus Slides. One section was stained with hematoxylin and eosin and scanned with a Keyence Fluorescence Microscope (BZ-X, Keyence). H&E images were used to guide microdissection of two subsequent sections.

Two sections were stained with hematoxylin and microdissected using an Arcturus XT LCM system onto CapSure HS LCM Caps (Thermo Fischer #LCM0215). Different well-defined groups of ducts were correlated with the clones identified by Atera CNV analysis, and mapped from the Atera-profiled slide onto the microdissection slides by an expert pathologist. Individual groups of ducts were targeted for microdissection, while avoiding the stroma between individual ducts. Normal breast ducts were separately dissected from each slide, while DCIS ducts from the same area were pooled between two slides.

### WGS data alignment and processing

Paired-end WGS reads from each of the 16 spatially dissected tumor regions were aligned to the human reference genome GRCh38 using BWA-MEM (bwa v0.7.19) with default parameters. Alignments were coordinate-sorted and indexed with SAMtools (v1.23.1). The complete workflow from the raw FASTQ through alignment and downstream copy number analysis was orchestrated with Snakemake (v9.23.1).

### WGS data copy number profiling

Copy number profiles were derived for each region with CNVkit (v0.9.13) in whole-genome mode. Tumor-only calling was performed against a flat reference with no control samples, such that log2 copy-number ratios are expressed relative to a uniform diploid expectation after correction for GC content, sequence repetitiveness and bin size. CNVkit automatically partitioned the accessible genome into bins, computed per-bin read depth, and generated bias-corrected per-bin log2 ratios. Ratios were segmented by circular binary segmentation, and integer copy number states were assigned from the segment log2 values assuming a diploid baseline. Segment log2 values were clipped to [−2, 2] for all downstream quantitative summaries and visualizations.

### Phylogenetic tree construction

For inferCNV results, all Leiden CNV clusters identified in copy number inference step were used in phylogenetic tree construction with all reference cells including Normal and Ductal Luminal (named as Merged Normal) as the root. For each clone, a mean genome-wide inferCNV profile was computed across all genomic bin regions. A region was scored as a gain (+) or loss (−) in a clone when at least 30% of that clone’s cells exceeded a per-cell region-mean magnitude of 0.02, with the diploid leaf fixed to the all-neutral state. With this clone-by-region character matrix, clonal phylogeny was reconstructed by exhaustive maximum parsimony using NumPy (v2.4.4): all unrooted binary topologies were enumerated and scored under an ordered (Sankoff) cost on the ordered states {−1, 0, +1} and the minimum-score topology was rooted on the Merged Normal so that evolution goes from normal to tumor. Branch lengths were set to the number of parsimony copy-number state changes along each branch. Ancestral states were reconstructed by Sankoff back-tracing and each branch was annotated with the discrete copy-number gains and losses acquired along it.

### Spatial niche assignment

Spatial niches were computed using *BuildAssayNiche* function (Seurat v5.5.0) using cell centroids while considering 30 nearest neighbors and 5 niches (neighbors.k = 30 and niches.k = 5). For this analysis, we used all annotated immune, stromal, and myoepithelial cells, and tumor epithelial luminal cells that had assigned CNV clones. Of note, this analysis included the Proliferating Tumor cell state cluster that was excluded from the CNV inference process.

### DCIS ducts segmentation

Tumor epithelial luminal cells were partitioned into ducts by single-linkage clustering: an undirected graph connected each cell to those of its 20 nearest tumor neighbors within 40 um, and connected components using function components() from the igraph (v2.3.2) package with at least 30 cells were retained as ducts (128 ducts, 21,587 of 24,505 tumor cells). Each duct was assigned its majority clone; median clonal purity 0.94 (19 ducts below 0.8). Duct niche-4 score is the mean over constituent cells.

### Quantifying immune niche fraction around DCIS clones

For each tumor cell, we defined a local microenvironment as all cells whose centroids lay within a 200 µm radius of the index cell, and scored it by the fraction of those cells assigned to the immune niche. Neighbors were retrieved by radius search (nn2, searchtype = “radius”, radius = 200; RANN v2.6.2), with the search cap set to k = 2,500 and verified to exceed the largest observed neighborhood so that no microenvironment was truncated. The index cell was excluded from both the numerator and the denominator, so the score is the number of immune-niche cells divided by the total number of other cells within 200 µm (median 310 neighboring cells, interquartile range 262–371). The denominator comprised cells of all types, making the score a local proportion that is insensitive to absolute cell density. Duct-level immune-niche fraction was then computed as the mean of the per-cell scores over the cells composing each duct.

### Quantification of the spatial co-occurrence of clonal stage and the immune niche

Association between duct clonal stage (Normal - Clone A - Clone B - Clone C - Clone D) and the fraction of the immune niche in a 200 µm radius around every duct was quantified by Spearman’s rho and Kendall’s tau; rank statistics were used because clonal stage is ordinal and immune niche fraction is strongly right-skewed. Because both quantities are spatially autocorrelated, neighboring ducts have similar clonal stages and similar immune-niche fractions; ducts are not independent observations, and the nominal P values returned by these tests are strongly anticonservative; they are therefore not reported. Significance was instead assessed by a torus-shift permutation test, which preserves the spatial structure of the immune-niche field while randomizing its position relative to the clonal field. In each permutation, the coordinates of all immune-niche cells were translated by a single uniformly random offset with wraparound (toroidal boundary) on the bounding rectangle of the section; every tumor cell’s local fraction was recomputed against this shifted pattern using its original neighbor count as the denominator, so that only the identity of cells as immune-niche or not was permuted and local cell density was left unchanged; duct means were re-formed and Spearman’s ρ recomputed. This rigidly translates the entire immune-niche point pattern, preserving its density, cluster sizes and autocorrelation, while destroying any real alignment with the clonal territories. One-sided P values, testing for positive co-localization, were computed as P = (1 + #{ρ_null ≥ ρ_obs}) / (1 + n_perm) over 999 permutations.

### Ordered-trend testing along the clonal phylogeny

Ducts were defined as above and each assigned its majority clone. For every duct, we computed the fraction of cells within 200 µm of its tumor cells that were SPP1+ TAMs and the fraction that were proliferating tumor cells. Monotone association of each readout with clonal advancement along the phylogeny order (normal - Clone A - Clone B - Clone C - Clone D) was tested with the Jonckheere–Terpstra statistic using the tie-corrected null variance. Significance was assessed by permutation: 999 permutations of duct clone labels and, for each 200 µm neighborhood metric, an additional 999 torus shifts in which the entire population point pattern was rigidly translated with wraparound on the section’s bounding rectangle and each duct score recomputed against its original neighbor count.

### Orion multiplex immunofluorescence

An unstained section from the DCIS FFPE block profiled with Atera was stained in two cycles as per the Rarecyte Immunofluorescence Staining of FFPE Tissue Samples for Orion^TM^ Analysis protocol and the Cyclic Immunofluorescence Staining of FFPE Tissue Samples for Orion^TM^ Analysis protocol. In brief, the section was baked overnight at 60 °C, deparaffinated and rehydrated. Antigen retrieval was performed for 5 minutes at 110 °C in Tris-EDTA. Autofluorescence was quenched while submerged in 4.5% H_2_O_2_/ 24nM NaOH in PBS under a light box for 60 minutes, followed by UV light for 30 minutes. The slide was then washed in Surfactant Wash Buffer (0.0025% Triton-X100 in PBS) and incubated in Image-iT FX Signal Enhancer (I36933, Invitrogen) for 15 minutes. The slide was washed in Surfactant Wash Buffer and stained for 2 hours using a multi-antibody panel in 5% mouse/5% rabbit serum (Supplementary Table 1). Twenty-two of these antibodies were purchased from Rarecyte, while nine were conjugated according to the Rarecyte Labeling Antibodies with ArgoFluor^TM^ Dyes protocol (CCR7, TCF1/7, KRT17, ISG15, KRT19, LAMC2, CLEC9A, KRT5, CD206). Next, the slide was washed in Surfactant Wash Buffer and stained for 30 minutes with Hoechst 33324 (H3570, Invitrogen) diluted 1:2000 in 10% normal goat serum. The slide was then mounted using ArgoFluor^TM^ Mounting Medium (Rarecyte, 42-1214-000) and #1.5 coverslips (VWR, 48393-231) and allowed to cure overnight. Whole slide scanning was performed using the Orion^TM^ scanner at 20X and images were extracted using an extraction matrix tuned to the antibody panel used. After scanning, the stained slide was decoverslipped after overnight incubation in PBS, followed by a 5 minute and 30 minute incubation in Piece^TM^ pH2 IgG Elution Buffer (ThermoScientific, 21028). The slide was then stained and scanned as described above, starting at the autofluorescence quenching stage.

### Cross-round image registration

The two 20-plex staining rounds of the same tissue section were co-registered into a single 40-channel image with ASHLAR v1.19.0, using the shared nuclear (Hoechst) channel as the alignment reference (maximum permitted shift, 30 µm). Round 2 was resampled onto the Round 1 coordinate frame, producing a full-resolution (0.325 µm pixel⁻¹) 40-channel pyramidal OME-TIFF.

## Data availability

All raw and processed WGS data for the laser-microdissected regions have been deposited to SRA under the BioProject ID PRJNA1509685. The multiplex IF data have been deposited using TissueViewer^29^ under this link: https://tissueviewer.com/link/m3zkg5.

## Code availability

The custom code for data preprocessing, analysis, and visualization is publicly available via Github at https://github.com/shaochwu/atera_project.

## Supplementary Figures

**Supplementary Fig 1.**
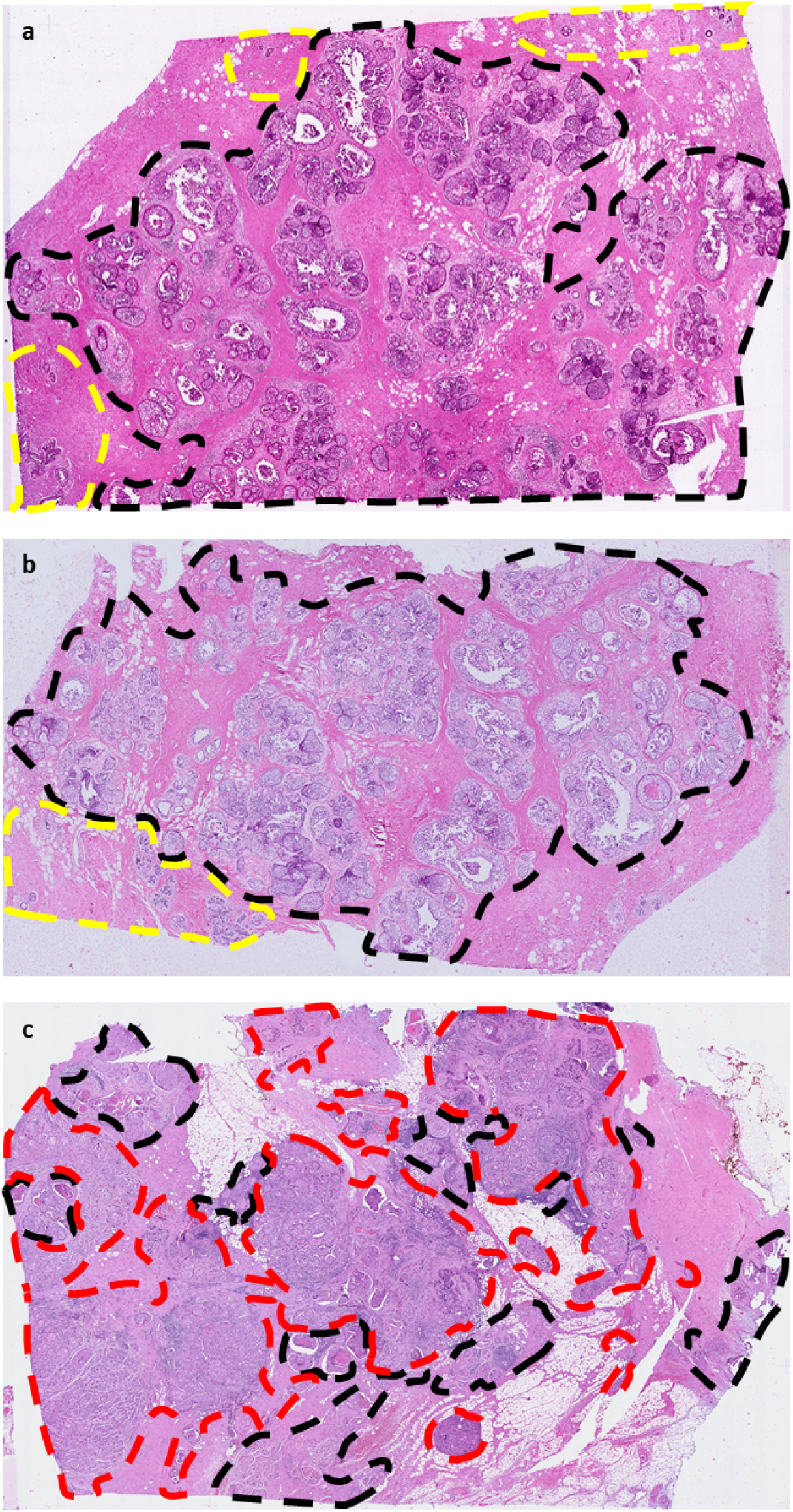
Histological overview of slides profiled by Xenium 280, Atera, and Xenium. Tissues profiled by Xenium 280 **(a)** and Atera **(b)** contain only DCIS (black) and normal breast glands (yellow), while the tissue profiled by Xenium 5K **(c)** contained DCIS (black) and invasive breast carcinoma (red).

**Supplementary Fig 2.**
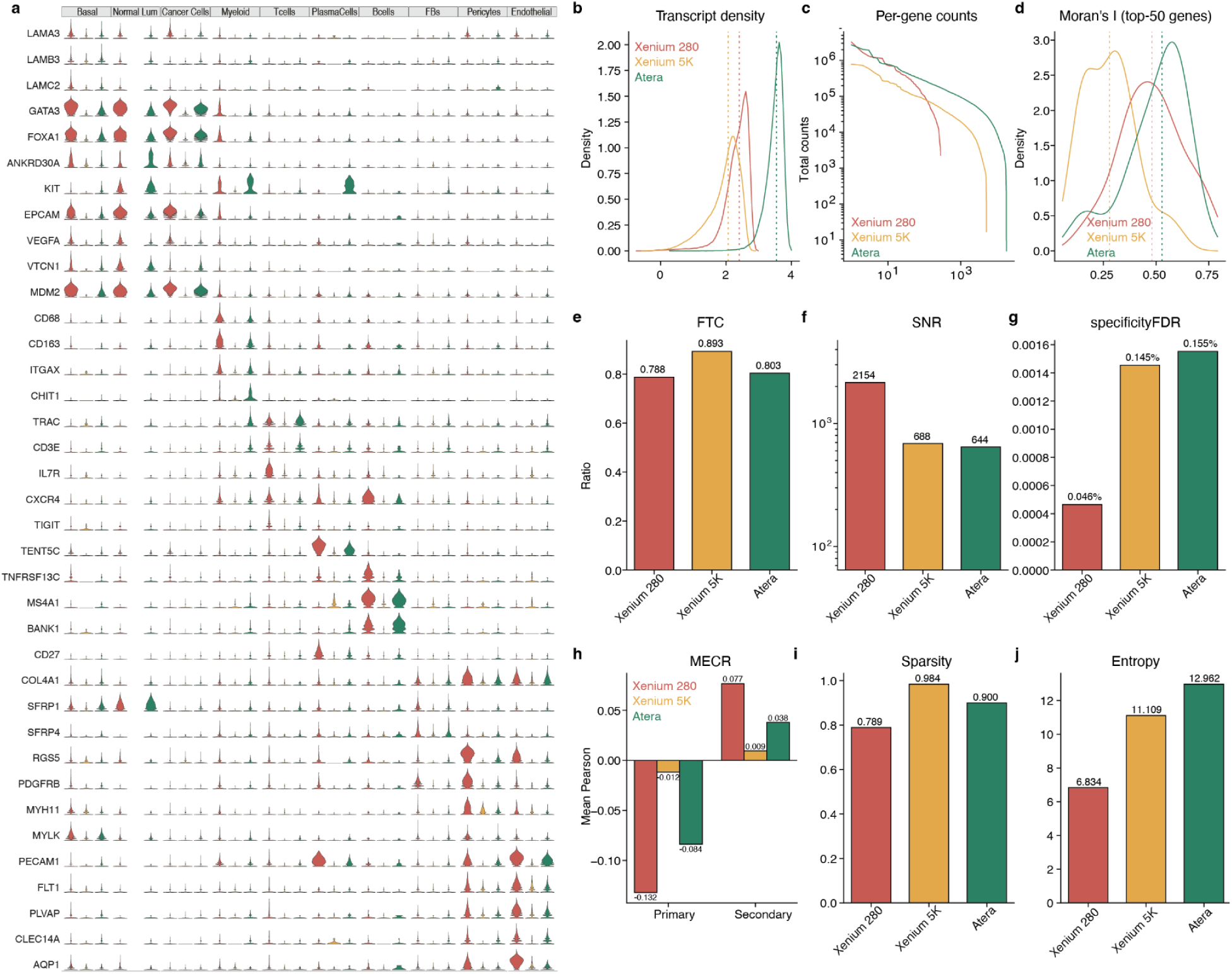
Marker-gene expression and SpatialQM quality-control metrics across three platforms. **(a)** Expression of the same marker genes across the annotated major cell types across the three platforms. **(b)** Distribution of transcript density (transcripts per 100 µm²). **(c)** Per-gene total counts across the three platforms. **(d)** Spatial autocorrelation (Moran’s I) for the top-50 genes. **(e)** Fraction of transcripts assigned to cells. **(f)** Signal-to-noise ratio (SNR). **(g)** False discovery rate (FDR) of how much of the detected signal is background rather than true transcripts. **(h)** Mutually-exclusive correlation ratio (MECR) for spatially distinct (‘primary’) versus co-localizing (‘secondary’) lineage pairs. **(i)** Sparsity across the three platforms. **(j)** Expression entropy across the three platforms.

**Supplementary Fig 3.**
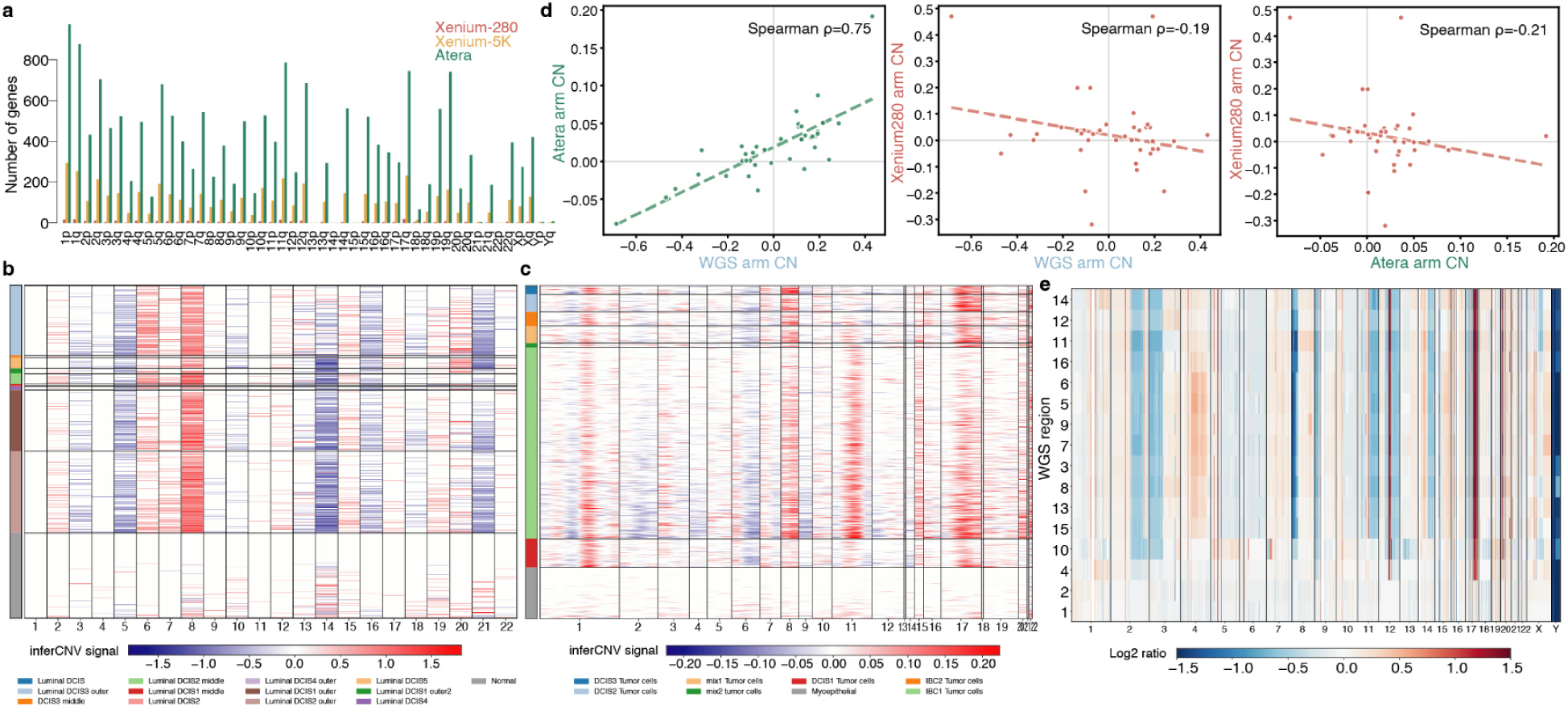
Chromosome-arm gene coverage, inferred copy number, and copy-number validation across platforms. **(a)** Number of genes detected per chromosome arm by Xenium 280 (red), Xenium 5K (orange) and Atera (green). **(b)** InferCNV heatmap for Xenium 280 (same DCIS block as Atera) with cells grouped by annotated luminal DCIS clusters. **(c)** InferCNV heatmap for Xenium 5K with cells grouped by annotated luminal DCIS and IBC clusters. **(d)** Arm-level copy-number correlation: Atera versus WGS (Spearman ρ = 0.75), Xenium 280 versus WGS (ρ = −0.19) and Xenium 280 versus Atera (ρ = −0.21); each point is one chromosome arm. **(e)** Copy number profiles of the 16 laser-microdissected regions by WGS.

**Supplementary Fig 4.**
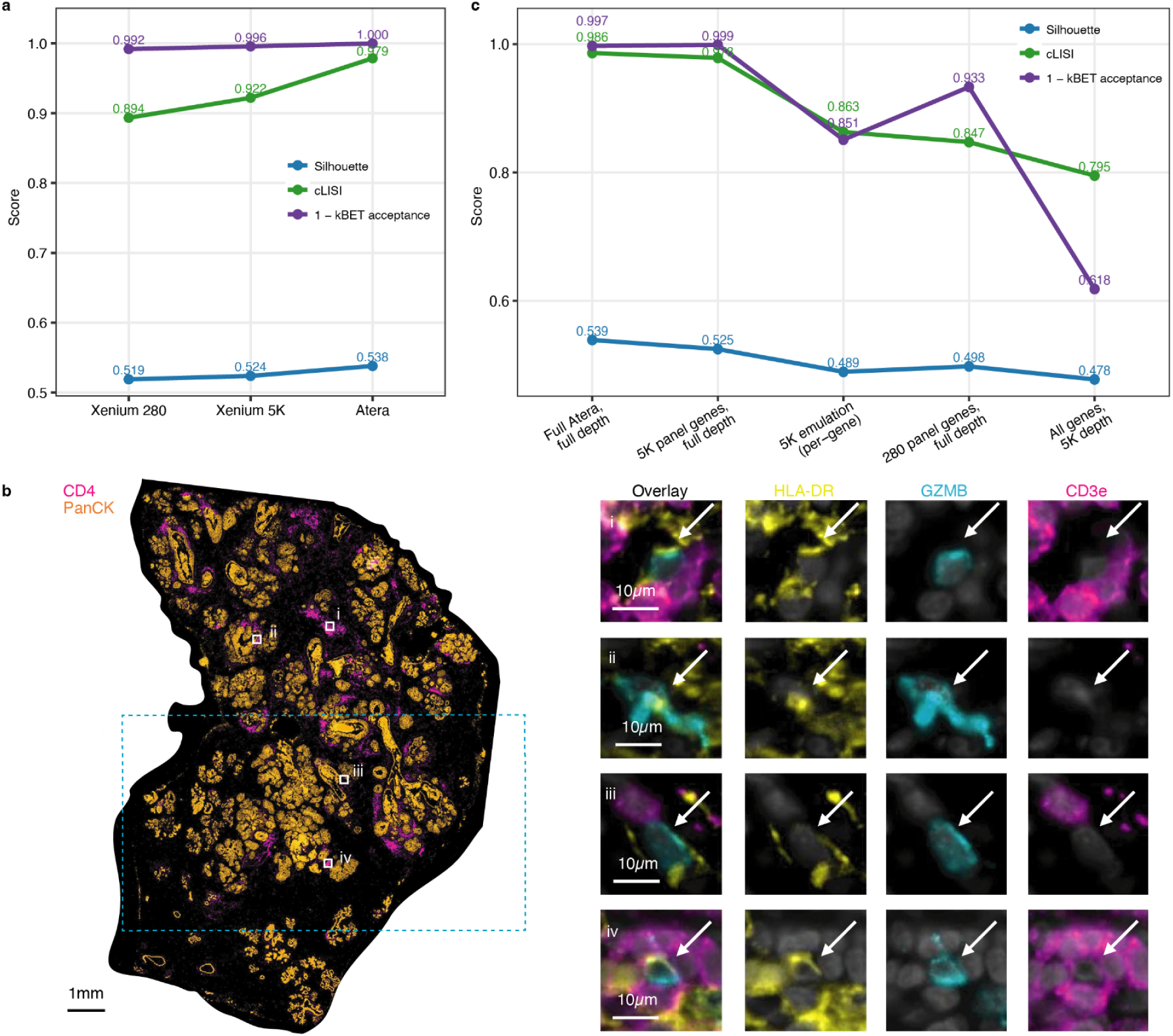
Resolution and validation of myeloid and rare immune populations across platforms. **(a)** Cluster-quality metrics for the whole reclustered myeloid compartment across Xenium 280, Xenium 5K and Atera. **(b)** Validation of rare cell types found through Atera by Orion multiplex IF on an adjacent tissue section. Left: whole section overview (CD4, Pan-CK) with the locations of the inserts. Right: i–iv show plasmacytoid dendritic cells, identified as HLA-DR and granzyme B positive cells that are negative for CD3e. **(c)** Myeloid annotation separability under reduced gene-panel and read-depth conditions; Atera myeloid cells were re-embedded from raw counts under five conditions in which only the gene set and/or per-cell depth varied.

